# HDgraphiX: A web-based tool for visualization of hydrogen deuterium exchange mass spectrometry data

**DOI:** 10.1101/2024.09.16.613363

**Authors:** Kent R. Vosper, Algirdas Velyvis, Siavash Vahidi

## Abstract

**Summary:** Hydrogen deuterium exchange mass spectrometry (HDX-MS) investigates protein structural changes by measuring deuterium incorporation into the protein amide backbone. Due to richness of information provided on protein conformational dynamics, HDX-MS data can be challenging to visualize effectively. To address this, we have developed HDgraphiX, a web-based tool that visualizes HDX data by processing outputs from two popular analysis software packages, DynamX (Waters Corp.) and HDExaminer (Sierra Analytics Inc). HDgraphiX performs statistical analyses, filters data based on statistical significance and presents the results in several forms of user-friendly publication-quality heatmaps (Chiclet plots). Unique features of this tool include the generation of Woods plots, volcano plots, and PyMOL colouring scripts, which are used to map deuterium uptake differences onto protein structures. Additionally, HDgraphiX offers numerous advanced options for customizing data processing and plotting without the need for manual data editing.

**Availability and Implementation:** HDgraphiX is available free of charge for all users at https://hdgraphix.net, the Python script and HTML template are deposited at https://github.com/KentV-UofG/HDgraphiX.

## Introduction

Hydrogen-deuterium exchange coupled to mass spectrometry (HDX-MS) is widely used in structural biology to detect changes in protein conformational dynamics in response to various stimuli.^1^ HDX involves the spontaneous exchange of labile hydrogen atoms in a protein for deuterium atoms in bulk solution.^2^ The HDX rate depends on the stability and solvent accessibility of hydrogen bonds at the exchange site.^3^ Continuous labeling HDX-MS, where deuterium uptake of the protein typically under two states (ligand-free vs ligand-bound) is measured and compared, is particularly popular in epitope mapping, monitoring binding interactions, and elucidating allosteric pathways.^4,5^ The HDX rate, and therefore deuterium uptake, often changes when ligand binds or conformation shifts, making accurate mass spectrometry-based measurement, rapid data extraction, and effective visualization of the data critical in the HDX-MS workflow.

HDX experiments generate complex and dense data from numerous partially overlapping peptides that require extensive processing. Software packages such as DynamX (Waters Corp.) and HDExaminer (Sierra Analytics Inc.), among others^6^, have significantly accelerated data analysis by efficiently extracting information from raw mass spectra and outputting them in a tabulated format. However, further processing is needed for statistical filtering and to create customizable publication-quality plots. Here, we introduce HDgraphiX, a web-based tool for the statistical filtering and visualization of HDX-MS data. While similar to existing HDX data processing and visualization tools like HD-eXplosion^7^, Deuteros^8^, HaDeX^9^, HDX-viewer^10^, HDXboxeR^11^ and MEMHDX^12^, HDgraphiX offers a broader range of plotting options and customization features for data processing and output. Additionally, HDgraphiX directly imports data from the commonly used DynamX and HDExaminer applications.

## Materials and Methods

HDgraphiX was written using Python 3.12 for data processing and formatting, while HTML, CSS, and JavaScript were used to format the website for ease of use. We used the following libraries within the Python code: NumPy (1.26.2), pandas (2.1.3), matplotlib (3.8.2), seaborn (0.13.0), and SciPy (1.12.0). Flask (3.0.2) was used to allow communication between the python script and HTML template for a user-friendly input.

## Results

HDgraphiX processes CSV outputs from two major HDX data analysis software packages, DynamX and HDExaminer, optionally performs a Welch’s *t*-test^13,14^, and generates three types of plots: Chiclet plots (with horizontal or vertical orientation), Woods plots, and volcano plots. A Chiclet plot is a 2-dimentional heatmap where the x-axis typically labels the peptides, and the y-axis represents D_2_O exposure time points. Differences in deuterium uptake between protein states are colour-coded in cells (Figure 1A). In a Woods plot, the x-axis shows the peptide position in the protein sequence, and the y-axis displays the difference in uptake. While Chiclet plots visualize differential deuterium uptake across multiple D_2_O exposure time points, Woods plots offer direct observation of peptide overlap and their positions in the sequence.^15^ Woods plots use many of the same plotting parameters as Chiclet plots (Figure 1B), but display the density of data differently, with separate graphs for each exposure time point rather than a single heatmap.^15^ Volcano plots show differential HDX data with the y-axis as -log_10_ transformed p-value from a Welch’s *t*-test and the x-axis as the difference in deuterium uptake between two states. Unlike the pass/fail statistical filtering applied to the Chiclet and Woods plots, volcano plots visualize the significance of uptake differences for each time point. Cutoff lines for the p-value and deuterium uptake significance are included for easy significance determination. HDgraphiX provides these plots for all possible permutations if more than two states are present or allows users to specify which states to compare.

**Figure 1:**
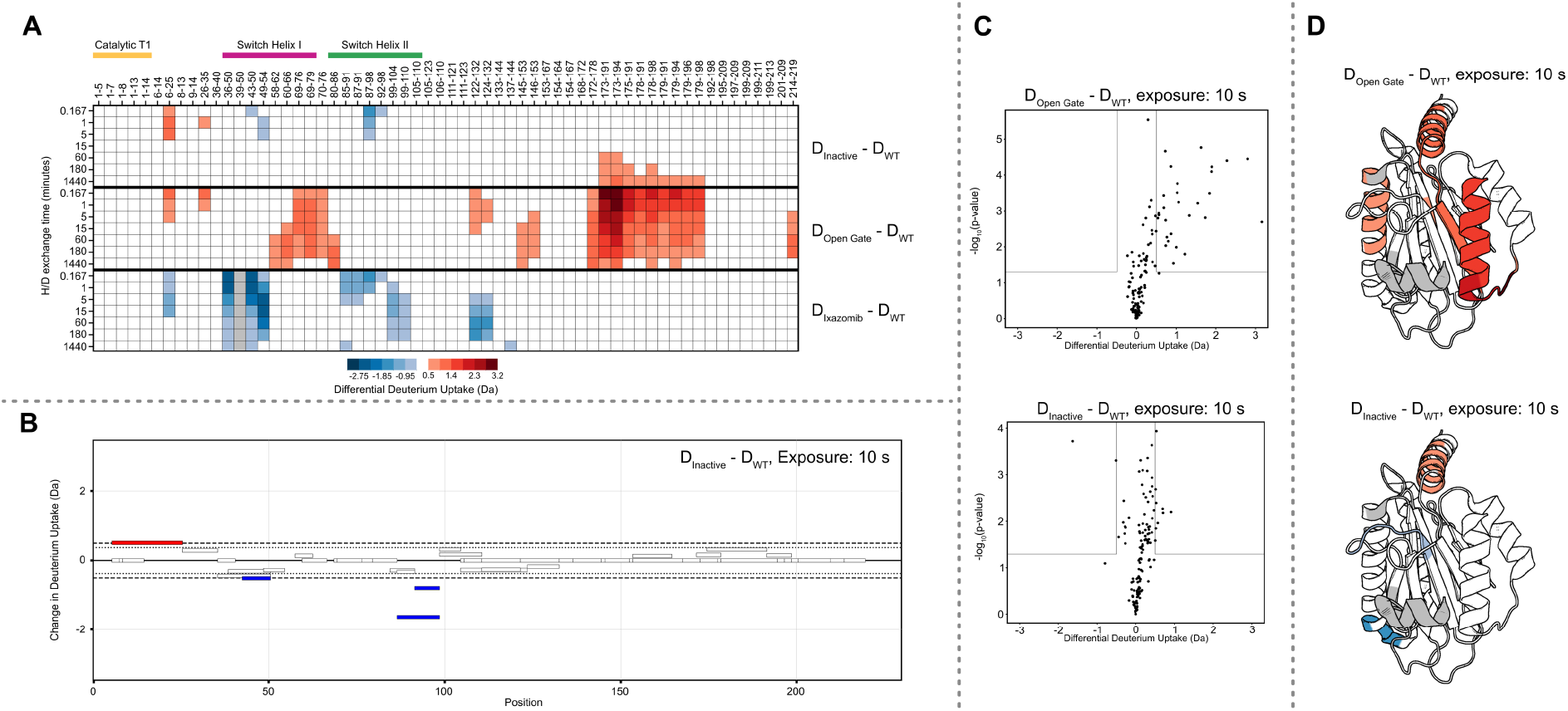
Output results of HDgraphiX. A) An example of a heatmap, or Chiclet plot, produced by HDgraphiX comparing three protein variants of the *M. tuberculosis* proteasome 20S core particle by Turner et al.^16^ The colour of each square indicates the difference in deuterium uptake between the compared states, according to the colour bar. For panels A, B, and D red represents an increase in deuterium uptake, and blue represents a decrease; B) An example of a Woods plot produced by HDgraphiX, examining differential D-uptake at the 10-second timepoint for the Inactive and WT variants. Peptides with differences in D-uptake beyond the set threshold are coloured; C) two examples of Volcano plots generated by HDgraphiX. The states protein being compared, and the deuterium exposure time are noted; and D) A comparison of the same timepoints as C, however, uses the heatmap colouring on the protein structure, visualized in PyMOL (PDBID 3MI0).

HDgraphiX’s versatility is its key advantage. Unlike existing data visualization tools, it allows for out-of-the-box use by automatically determining the number of replicates, and HDX data bounds for rapid plot generation. Users can opt for the simplest setting, where the code determines the maximum differential deuterium uptake value in the dataset, sets the differential deuterium uptake range based on this computed maximal value, and plots the input data using a standard colour palette with shades of red for increased deuterium uptake and shades of blue for decreased uptake. Users simply upload a CSV output from DynamX or HDExaminer and click “Generate Plot”. Further bound and colour options offer granular control over the plots including manual range settings, alternate colour schemes, or inputting hexadecimal colour values for full customization. HDgraphiX offers additional customizations such as i) renumbering peptides for consistency with previously published work; ii) removing unwanted deuterium exposure times or peptides; iii) using a mutation dictionary to compare the deuterium uptake differences across peptides with amino acid substitutions; and iv) precise domain labeling with user-defined start and end points. A full description of the customization options is available in the help document, which can be downloaded from the website. The plot outputs include a series of PNG files or a series of PDF files containing plots in vector format. HDgraphiX also outputs PyMOL coloring scripts that color a PDB structure according to differential deuterium uptake, based on the generated heatmap, thereby aiding in the visualization of structural changes within the context of the protein structure (Figure 1D).

## Acknowledgements

We thank the members of the Vahidi lab for their assistance in testing HDgraphiX.

## Data availability

No new data were generated or analysed in support of this research.

